# Harnessing cytoplasmic particles movement of the human early embryo analysed by advanced imaging and artificial intelligence to predict development to blastocyst stage

**DOI:** 10.1101/2020.11.26.399832

**Authors:** Giovanni Coticchio, Giulia Fiorentino, Giovanna Nicora, Raffaella Sciajno, Federica Cavalera, Riccardo Bellazzi, Silvia Garagna, Andrea Borini, Maurizio Zuccotti

**Author notes:** Contributed equally to the manuscript (Giovanni Coticchio, Giulia Fiorentino). **Correspondence** Giovanni Coticchio: 9.baby Family and Fertility Center, Via Dante, 15, 40125 - Bologna, Italy, Maurizio Zuccotti: Laboratory of Developmental Biology, Department of Biology and Biotechnology ‘Lazzaro Spallanzani’, University of Pavia, Via Ferrata, 9, 27100 – Pavia, Italy.

## Abstract

**Research Question:** Progress in artificial intelligence (AI) and advanced image analysis offers unique opportunities to develop novel embryo assessment approaches. In this study, we tested the hypothesis that such technologies can extract and harness novel information derived from cytoplasmic movements of the early human embryo to predict development to blastocyst.

**Design:** In a proof-of principle study, an artificial neural network (ANN) approach was undertaken to assess retrospectively 230 human preimplantation embryos. After ICSI, embryos were subjected to time-lapse monitoring for 44 hours. For comparison as a standard embryo assessment methodology, a single senior embryologist assessed each embryo to predict development to blastocyst stage (BL) based on a single picture frame taken at 42 hours of development. In the experimental approach, in embryos that developed to blastocyst or destined to arrest (NoBL), cytoplasm movement velocity (CMV) was recorded by time-lapse monitoring during the first 44 hours of culture and analysed with a Particle Image Velocimetry (PIV) algorithm to extract quantitative information. Three main AI approaches, the k-Nearest Neighbor (k-NN), the Long-Short Term Memory Neural Network (LSTM-NN) and the hybrid ensemble classifier (HyEC) were employed to classify the two embryo classes.

**Results:** Blind operator assessment classified each embryo in terms of ability of development to blastocyst, reaching a 75.4% accuracy, 76.5% sensitivity, 74.3% specificity, 74.3% precision and 75.4% F1 score. After integration of results from AI models together with the blind operator classification, the performance metrics improved significantly, with a 82.6% accuracy, 79.4% sensitivity, 85.7% specificity, 84.4% precision and 81.8% F1 score.

**Conclusions:** The present study suggests the possibility to predict human blastocyst development at early cleavage stages by detection of CMV and AI analysis. This indicates the importance of the dynamics of the cytoplasm as a novel and valuable source of data to assess embryo viability.

## Introduction

One of the long-term, and among the most important, goals of medically assisted reproduction (MAR) is the achievement, in the shortest possible period of time, of a single healthy term pregnancy following the transfer of a single embryo. To this end, embryos need to be assessed by non- or minimally-invasive methodologies and prioritized for transfer according to their developmental potential. Until the early 2000’s, in the vast majority of MAR programs, embryos were being cultured until day two or three post-insemination and selected for transfer according to static morphological criteria. More recently, significant paradigm shifts have emerged, based on the extension of embryo culture to day five or six to discriminate, by self-selection, embryos able to develop to the blastocyst stage. These blastocysts, on average with higher developmental potential compared with cleavage-stage embryos, can be used for embryo transfer with increased chances to achieve a viable pregnancy per transfer attempt (Glujovsky et al., 2016). To the aim of further increasing efficiency, i.e. to identify more precisely and prioritize for transfer the most viable embryo in a cohort, blastocysts can be assessed according to static (“The Istanbul consensus workshop on embryo assessment: proceedings of an expert meeting.”, 2011) or dynamic (Gallego et al., 2019) morphological criteria, or even by adopting a strategy of preimplantation genetic screening for aneuploidies (PGT-A) (Griffin and Ogur, 2018). Collectively, indeed, these approaches based on blastocyst culture assure a higher treatment efficiency compared with day two or three embryo transfer. However, they cannot be considered ideal for several reasons. A major drawback of such methodologies lies in the culture to the blastocyst stage *per se*. In fact, blastocyst culture is more expensive and more demanding for the management of the IVF laboratory workflow. Even more importantly, if not mastered properly, blastocyst culture may impact on intrinsic embryo developmental potential, affecting overall treatment efficacy (Swain, 2019). This scenario clearly points towards the need for a more advanced, non-invasive embryo assessment methodology able to predict developmental potential at earlier stages, without necessarily resorting to extended culture.

Progress in the realm of artificial intelligence and in image analysis offers unique opportunities to develop novel embryo assessment approaches. Recently, promising attempts were made to harness the potential of deep learning to predict embryo implantation potential (Khosravi et al., 2019; Miyagi et al., 2019; Tran et al., 2019). However, all such cases involved the use of static images or videos of blastocyst stage embryos, leaving the question of prediction at earlier stages unanswered.

In this proof-of-principle study, we approached the option of advanced image analysis of early development to predict ability of blastocyst formation. To this aim, sequential images collected by bright-field time lapse microscopy (TLM) over less than two days of culture were subjected to particle image velocimetry (PIV) analysis to detect the dynamics of cytoplasmic movements. The resulting data were elaborated by neural network-based artificial intelligence (AI). The strategy to investigate cytoplasm dynamics as a novel source of data to assess oocyte function and embryo viability finds justification in studies carried out in the mouse model (Ajduk et al., 2011;Yi et al., 2013; Bui et al., 2017). Nevertheless, movement of cytoplasmic particles has been until now totally neglected in attempts to predict human embryo viability.

The study results suggest that three powerful technological and analytical tools, TLM, PIV and AI, can be combined to extract predictive information on human embryo viability from cytoplasm dynamics occurring at early developmental stages.

## Materials and Methods

This retrospective proof-of-principle study included 230 embryos generated in ICSI cycles carried out between October 2015 and May 2018. Approval for the study was obtained from the local Institutional Review Board. Laboratory non-clinical data were used for research purposes only. Cases suitable for analysis were selected according to the rule to have two sibling embryos from the same cohort (cycle), of which one developed to blastocyst stage and one arrested to an earlier stage. In this fashion, we aimed at limiting possible biases due to patient variability.

Diagnosis of infertility included various causes, including male factor, tubal factor and polycystic ovary (PCO, but not PCOS) with or without chronic anovulation. Such inclusion criteria were chosen because consistent with the expectation to have a relatively high number of embryos in a cohort and the ensuing possibility to extend embryo culture to the blastocyst stage, a condition required by the study design. Ovarian stimulation was carried out as previously described by Zacà et al. (Zacà et al., 2018).

### Semen preparation and ICSI procedures

Semen selection for ICSI was performed by discontinuous PureSperm (Nidacon, Gothemberg, Sweden) gradient (Borini et al., 2006). After preparation, sperm were evaluated for concentration, total and progressive motility and morphology, according to WHO procedures (World Health Organization, 2010).

### Embryo culture and time lapse image acquisition

Oocytes/embryo culture was carried out using the EmbryoScope equipment (Vitrolife, Goteborg, Sweden), an integrated time-lapse technology-incubator system for embryo-culture, carried out in a N2/CO2/O2 (89:6:5, v/v) atmosphere at 37°C without control of humidity. Microinjected oocytes were placed inside pre-equilibrated slides (EmbryoSlide; Vitrolife), each containing 12 droplets of 25 μl of cleavage medium (Cook IVF, Australia) covered by 1.2 ml of mineral oil (SAGE, Biocare Europe, Rome, Italy). On day 3, cleavage medium was replaced with blastocyst medium (Cook IVF, Australia) to extend embryo culture until day 5, where appropriate (Lagalla 2017). Images were acquired over a period of 125 hours starting from the time of ICSI, with a 15 min interval between consecutive picture frames.

### Operator-based prediction of development to blastocyst stage

The outcome of AI analyis was first comparatively assessed and then combined with standard operator-based prediction of development to blastocyst. Embryo assessment by standard morphology was preferred to alternative assessement approaches, i,e, morphokinetic algorithms, because static evaluation is routinely performed following the recommendations of an international consensus ((“The Istanbul consensus workshop on embryo assessment: proceedings of an expert meeting.”, 2011) and remains the “standard of care” in clinical embryology. On the other hand, while having great potential, morphokinetic evaluation has not reached yet a similar degree of consensus, perhaps also as an effect of biases derived from different technological platforms and applicative protocols. On day two, at 42 hours post-ICSI a single embryologist with over 20 years of experience in clinical embryology assessed each embryo according to: a) blastomere number, size and mutual position; b) degree (percentage of total volume) of fragmentation; c) multinucleated blastomeres; d) cellular dysmorphisms (e.g., large vacuoles) (“The Istanbul consensus workshop on embryo assessment: proceedings of an expert meeting.”, 2011). Based on such morphological assessment performed statically (i.e., at a single time point), the operator formulated a binary prediction (yes/no) of development to blastocyst stage according to the recommendations of the Alpha and ESHRE Istanbul Consensus (“The Istanbul consensus workshop on embryo assessment: proceedings of an expert meeting.”, 2011).

### Definition of blastocyst

At 116 hours post ICSI embryos were categorized as developed at the blastocyst stage if they showed at least the following minimal requirements: a) at least a Grade 1 blastocoel expansion; b) a discernible cluster of cells forming the inner cell mass (ICM); c) a wall of epithelial-like cells delimiting the blastocoel and enclosing the ICM (“The Istanbul consensus workshop on embryo assessment: proceedings of an expert meeting.”, 2011).

### Cell_PIV analysis

Cell_PIV software (kindly provided by Dr. Shane Windsor), based on the Particle Image Velocimetry (PIV) algorithm, was used to detect embryo movement velocity (EMV) during the first two-cell division (44 hours) of BL and No-BL embryos. The movement velocity vectors, observed in the embryo, were calculated by cross correlation between the patterns of the pixels of adjacent video frames, extracting five features: *(i) Direction* (D), the mean of the inverse of the local standard deviation of the direction of the vectors; *(ii) Vorticity* (V), the mean vorticity of the vectors in each frame; *(iii) Hybrid* (H), a hybrid combination of the mean magnitude and inverse standard deviation data; *(iv) Meanmag* (M), the mean magnitude of the vectors in each frame; *(v) Summed* (S), the mean magnitude of the vectors summed over a set number of frames forward in time from the current point.

### Statistics

Differences in BL and No-BL embryos cytoplasmic movements were evaluated using the Two-sample Kolmogorov-Smirnov test with the MATLAB software (R2018B). Data were considered significantly different when p < 0.05.

### Artificial Intelligence

Both k-NN and LSTM-NN were implemented with MATLAB software (R2018B). In particular, the LSTM-NN was developed using the MATLAB Deep Learning Toolbox. A set of k-NN and LSTM-NN models, with empirically established parameters and different combination of features, was defined. Among them, four models (k-NN-1, k-NN-2, LSTM-NN-1, LSTM-NN-2) used for final comparison with the operator on the test set, were chosen based on the best results of the cross validation on the training set. The selected models were then trained on the entire training set. To evaluate the models generalization capabilities, a 10-fold cross validation on the entire dataset (230 embryos, 118 in NoBL class and 112 in BL class) was also performed for the k-NN-1, k-NN2, LSTM-NN-1 and LSTM-NN-2 algorithms (Figure 5). For the k-NN models, we set the number of neighbours equal to four and the similarity between embryos was computed coupling the Dynamic Time Warping with the Kullback distance. The LSTM-NN, whose architecture is shown in Figure 3, was trained with the Adam optimization function for 100 epochs, with a learning rate of 0.1 and a batch size of 20. Results of the AI models on the test set were evaluated in terms of accuracy (BL-NoBL recognition rate), sensitivity (True-BL recognition rate) and specificity (True-NoBL recognition rate).

## Results

Immediately after ICSI, each of the obtained 230 zygotes was transferred in a single Embryoscope well and cultured for a total of 125 hours (5.2 days). Of these, 112 reached the blastocyst stage (48.7%), whereas 118 arrested sometime earlier, but never before the second cleavage cycle.

Based on a single picture frame taken at 42 hours culture post ICSI, an expert operator blindly classified each of the 230 embryos - according to previously established morphological criteria (“Istanbul consensus workshop on embryo assessment: proceedings of an expert meeting.”, 2011) - as developmentally competent (BL) or incompetent (NoBL), reaching a 75.4% accuracy, 76.5% sensitivity and 74.3% specificity.

The time-lapse sequence recorded during 44 hours (175 frames), corresponding to completion of the second embryonic cell cycle division, was analysed using Particle Image Velocimetry software Cell_PIV (Figure 1A), with five different image processing features: *Direction*, *Vorticity*, *Hybrid*,

**Figure 1.**
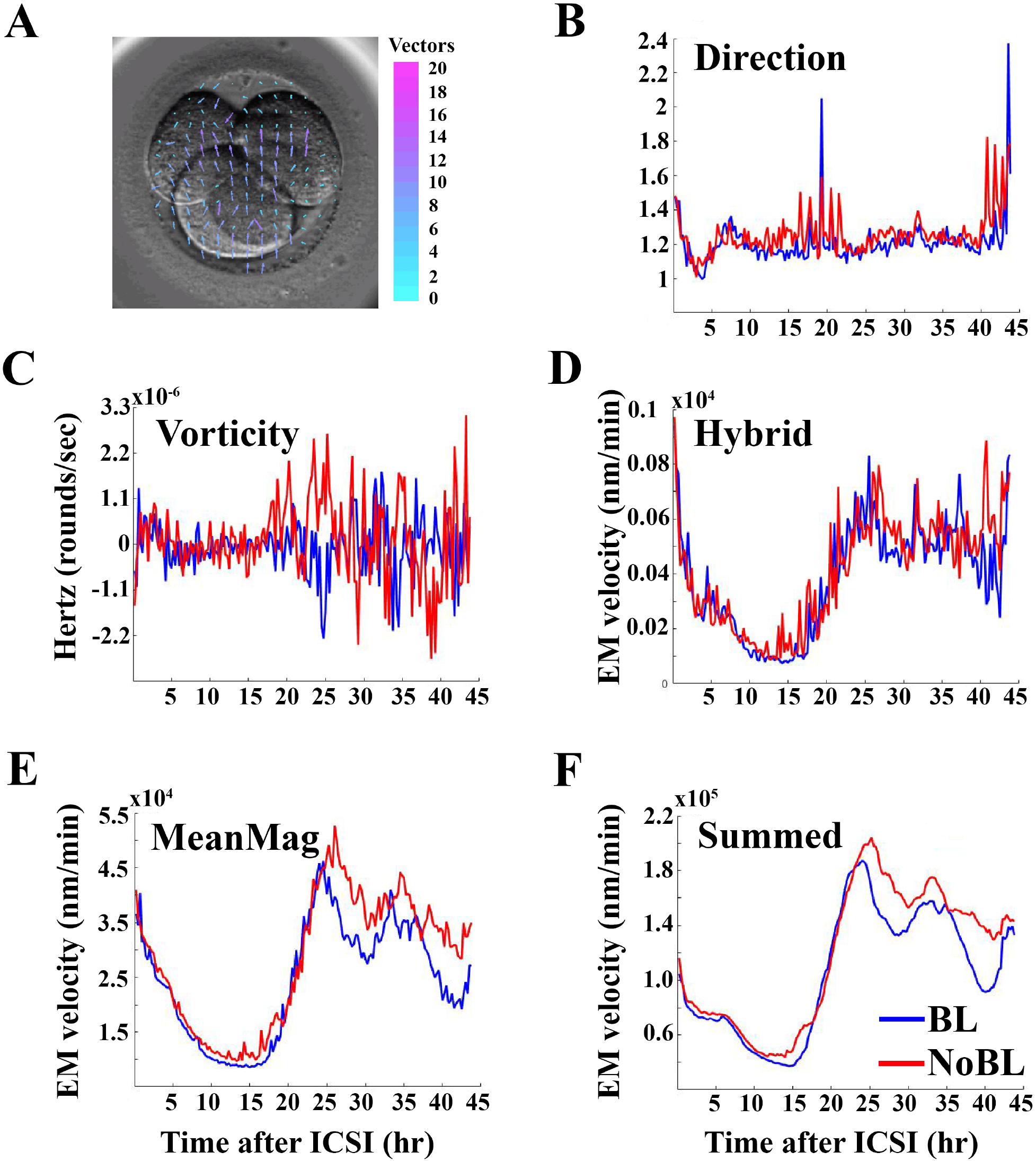
Embryo movement velocity profiles of BL and NoBL embryos. (A) Representative image frame of an embryo at 32 hours after ICSI showing the Cell-PIV velocity vectors. The color and length of the arrows (vectors) indicate the velocity module of the movements when comparing the previous frame with that pictured. Colored vector scale bar: blue, lower velocity; purple, higher velocity. (B-F) EMV profiles analysed using *Direction, Vorticity, Hybrid, Meanmag* or *Summed* Cell_PIV feature, respectively.

### *Meanmag* and *Summed* (see M&M)

Analysis of the first three parameters highlighted noisy and overlapping profiles of BL and NoBL embryo movement velocity (EMV) (Figure 1B-D); instead, the *Meanmag* (Figure 1E) and, more significantly, the *Summed* (Figure 1F) features identified two main and consistent time intervals with statistically different EMVs.

Interestingly, when the *Summed* EMV temporal profiles of BL and NoBL embryos were correlated with known key events occurring during the first 44 hours of development (Figure 2), the first time-interval overlapped with pronuclear re-positioning and ended shortly before the time of the first embryonic cell division. The second time-interval coincided with the second cleavage cycle, until the end of the time-lapse frames analysed.

**Figure 2.**
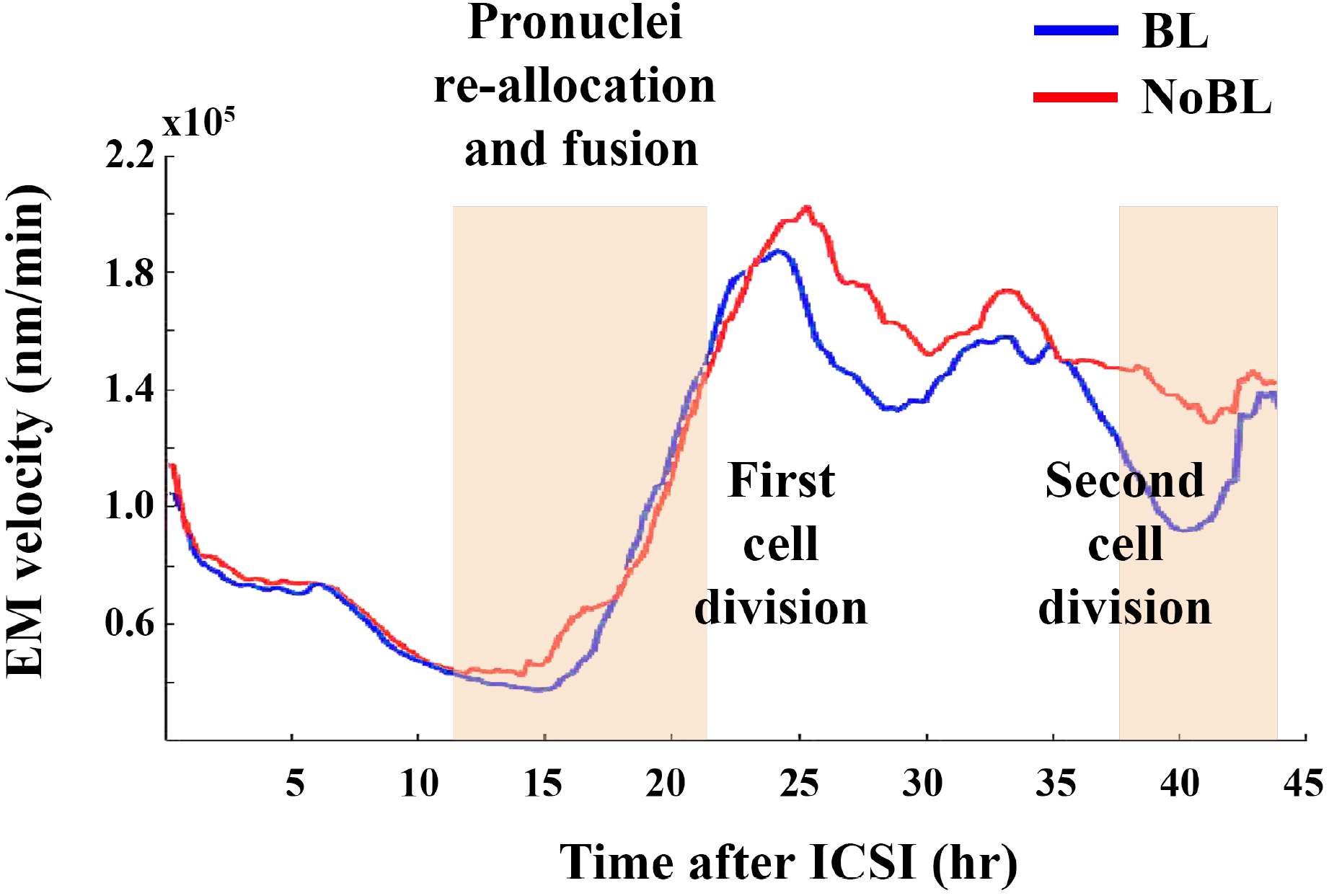
Cytological events occurring during the first two segmentation divisions. The *Summed* EMV profile of BL and NoBL embryos resulted significantly different during the time-intervals corresponding to pronuclei re-allocation and the second embryonic division.

Whilst the morphodynamic patterns measured by Cell_PIV suggest, at least at these two time-intervals, the potential to discriminate between the two embryo classes, their high cytoplasm movement velocity (CMV) variability restricts their possible use for a single embryo classification in the clinical practice. Nevertheless, this limitation was resolved by the subsequent AI analysis.

To this end, here we used two AI approaches, the k-Nearest Neighbor (k-NN) and the Long-Short Term Memory Neural Network (LSTM-NN), whose algorithms have been exploited earlier in different biomedical classification contexts to reveal hidden patterns from complex biological data and to support the clinical decision process (He et al., 2019).

Out of 230 embryos analysed, 161 (70%) were randomly selected as training set of the AI algorithms, whereas the remaining 69 (30%) were employed as test set. Both algorithms were fed with either the data of a single Cell_PIV feature (i.e., M or S) or with a combination (i.e., M+S or D+V+H+M+S).

### k-NN

The k-NN algorithm, largely used to solve time series classification problems (Yakovitz, 1987), classifies a new sample based on its distance similarity with the training set. Since the computation of the distances between two time series is frequently subject to slight time shifts, the k-NN on Cell_PIV temporal series was preceded by the application of Dynamic Time Warping (DTW), a pre-processing technique that produces the optimal alignment between sequences (Chaovalitwongse et al., 2007; Müller et al., 2007; Geler et al., 2016; 2020; Tran et al., 2019).

Of the four k-NN implementations performed, the D+V+H+M+S (k-NN-2) showed the highest accuracy (72.5%), the best balance between sensitivity (76.5%) and specificity (68.6%), the highest precision (70.3%), and a F1 score equal to 73.2%; instead, the M+S (k-NN-1) approach gave a similar accuracy (71.0%), lower specificity (60.0%) and precision (66.7%), but a higher sensitivity (82.3%) and F1 score (73.7%). Implementation with a single feature (i.e., M or S) resulted in low accuracy (< 65%) and, thus, this approach was abandoned.

### LSTM-NN

LSTM-NN is a non-linear computational approach able to encode and store temporal information in its layers and, through subsequent nodes, convert it into a classification output (Figure 3). For this reason, LSTMs have been widely used to classify time series data (Karim et al., 2018). When trained on M+S features (LSTM-NN-1), LSTM-NN showed a 71.0% accuracy, 76.5% sensitivity, 65.7% specificity, 68.4% precision and 72.2% F1 score; instead, the D+V+H+M+S combination (LSTM-NN-2) displayed the same accuracy (71.0%), lower sensitivity (61.8%) higher specificity (80.0%), 75.0% precision and 67.7% F1 score. Implementation with a single feature (i.e., M or S) resulted in low accuracy (< 70%) and, as before, this approach was abandoned.

**Figure 3.**
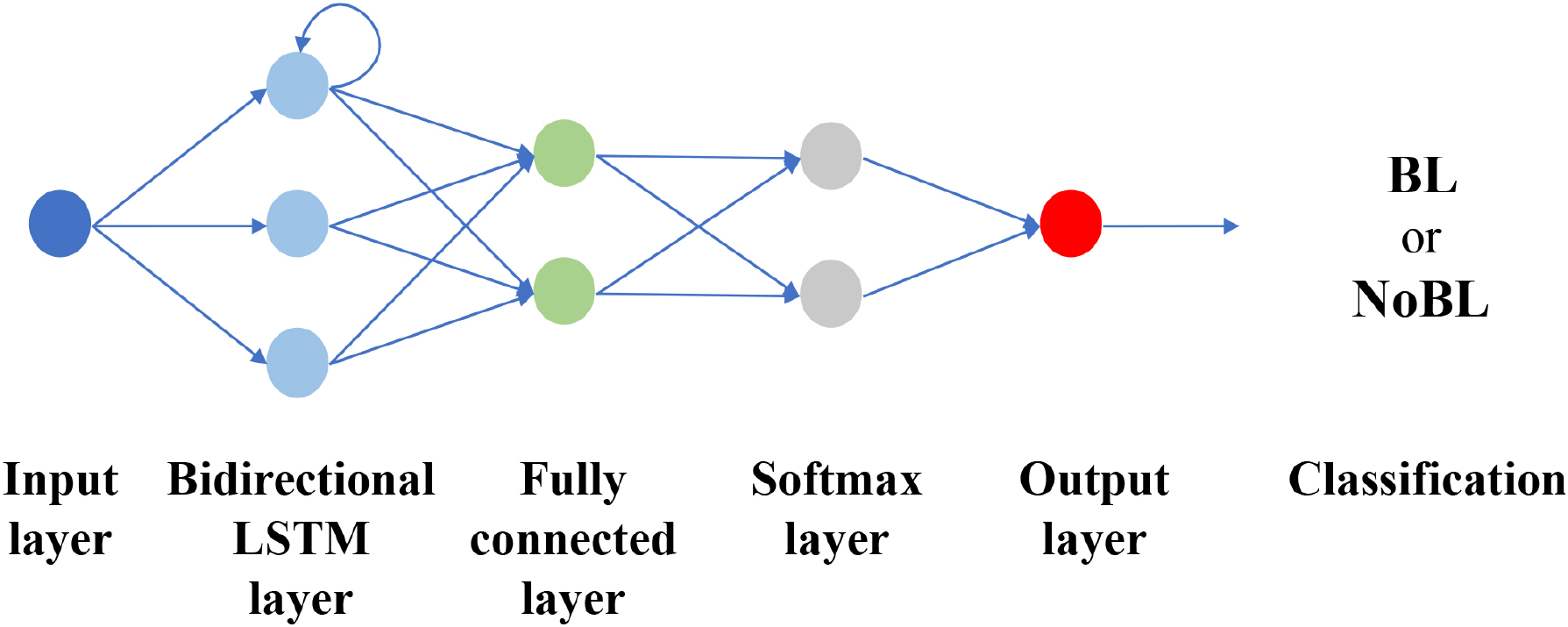
LSTM Neural Network architecture. The structure of the LSTM-NN is made of five main layers. An input layer, whose number of nodes is equal to the number of features used; a bidirectional LSTM layer, made with 3 nodes, whose function is to bring out hidden patterns along the time series; a fully connected layer and a softmax layer, convert the temporal information stored in the LSTM layer in a classification probability; an output layer provides the class with the highest probability.

### Hybrid ensemble classifier

The results of the AI models (k-NN-1, k-NN-2, LSTM-NN-1 and LSTM-NN-2), showed a similar classification accuracy to that reported by the operator (75.4%), although their comparison highlighted a high classification variability for each single embryo (Figure 4A).

**Figure 4.**
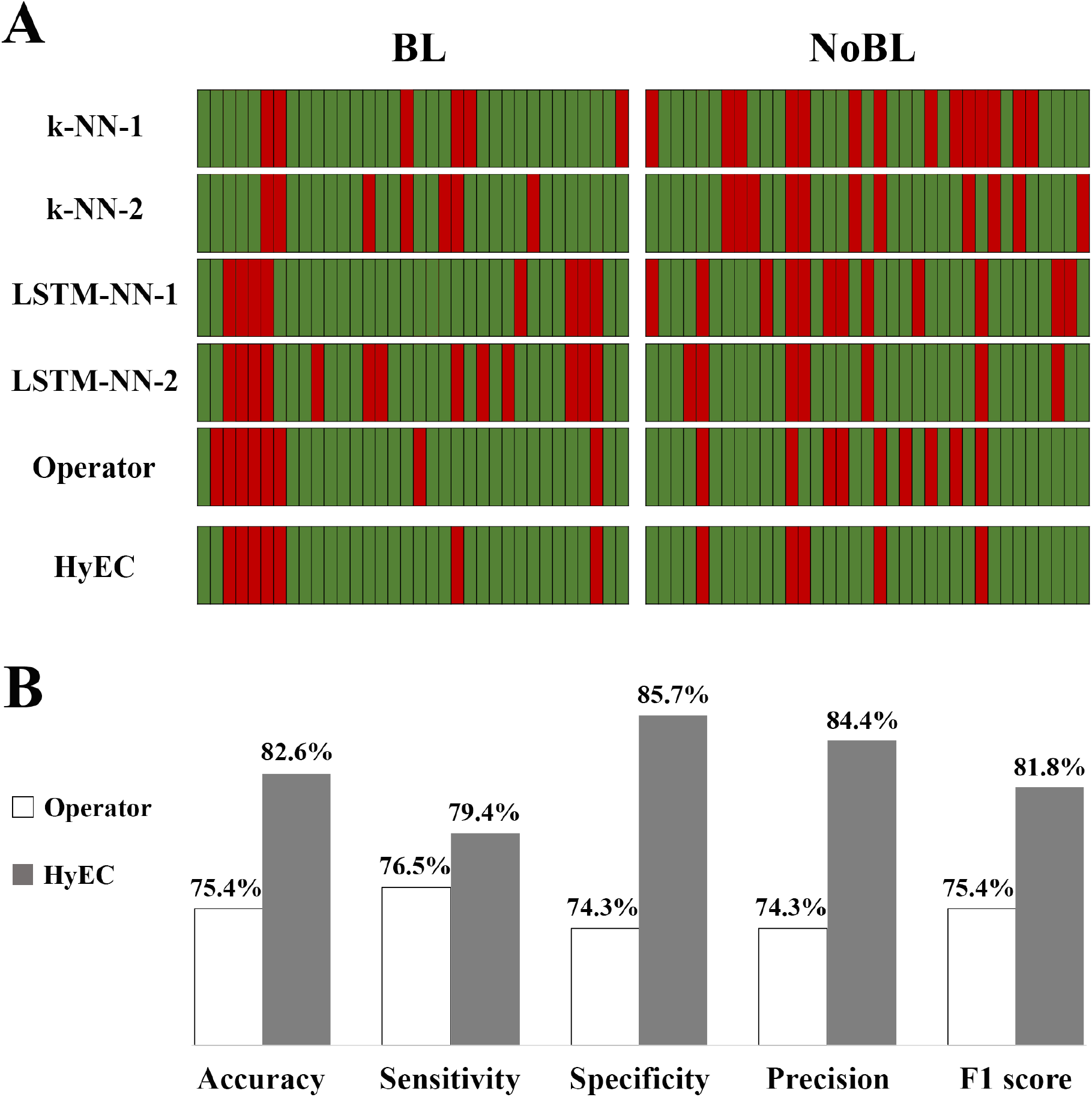
Improved embryo-quality classification with *hybrid ensemble classifier*. (A) Test embryos correctly (green) or incorrectly (red) classified by the AI models (k-NN-1, k-NN-2, LSTM-NN-1 or LSTM-NN-2), by the operator or the *hybrid ensemble classifier* (HyEC). (B) Performance metrics obtained by the operator alone or by the HyEC.

Next, we employed an ensemble classifier to integrate the results of the four AI models together with the blind classification made by the expert operator, assuming, as decision rule, the agreement of at least 3 out of 5 classifiers. For its characteristics, this classifier was named *hybrid ensemble classifier* (HyEC). The adoption of this strategy led to an improvement of all the performance metrics, with a final 82.6% accuracy, 79.4% sensitivity, 85.7% specificity, 84.4% precision and 81.8% F1 score (Figure 4B).

In addition, the generalization capability of each of the four models and of the HyEC was evaluated by 10-fold cross validation on the entire dataset (230 embryos, 118 in NoBL class and 112 in BL class) (Figure 5).

**Figure 5.**
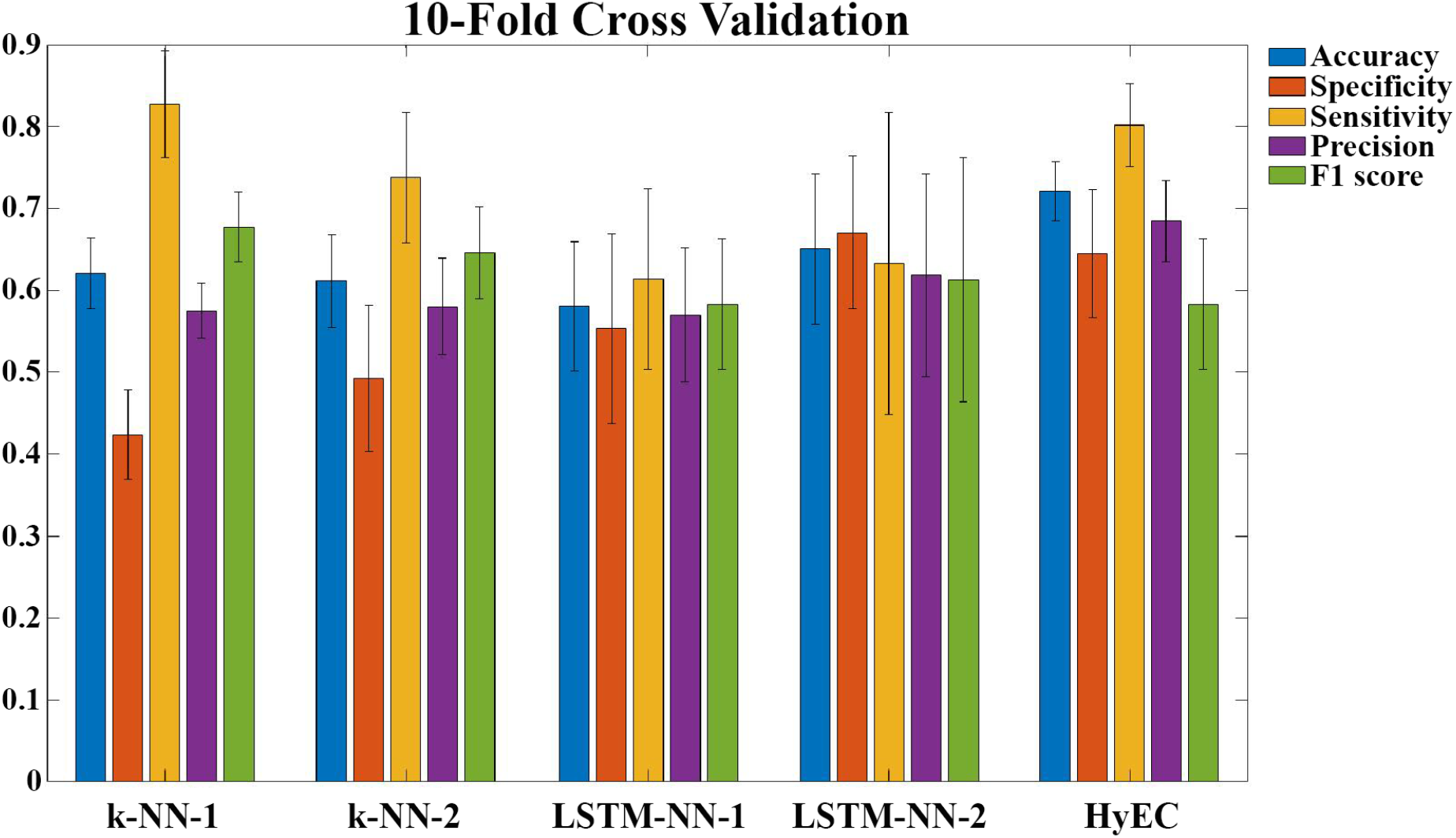
10-fold cross validation Mean and 95% Confidence Intervals of Accuracy, Specificity, Sensitivity, Precision and F1 score, computed on the entire dataset (230 embryos, 118 in NoBL class and 112 in BL class) for the k-NN-1, k-NN-2, LSTM-NN-1, LSTM-NN-2 and HyEC classifiers.

## Discussion

The present study was designed to explore the use of a novel source of data to predict the ability of the human embryo to develop to the blastocyst stage. This aim was pursued by harmonizing different methodologies, i.e. TLM, advanced image analysis and AI computational and elaboration tools. Collectively, the resulting data indicate that our approach can achieve a diagnostic accuracy classifiable as “very good” (Šimundić, 2009). This is noteworthy, considering the proof-of-principle character of our study and the small size of the data set analysed. However, such findings should be interpreted only in a research context, lacking at present validation of their clinical applicability.

The current standard of embryo assessment in MAR relies on static morphological parameters (“The Istanbul consensus workshop on embryo assessment: proceedings of an expert meeting.”, 2011). In the large majority of clinics worldwide, embryos are ranked according to their presumed developmental potential and prioritized for transfer at the cleavage (day three) or blastocyst (day five-six) stage. While the two approaches generate comparable cumulative live birth rates after the transfer of fresh and cryopreserved embryos (Glujovsky et al., 2016), self-selection derived from blastocyst culture offers the advantage of a shorter time to pregnancy by reducing the number of embryos suitable for transfer. However, because blastocyst culture is not immune from technical and management drawbacks, early and robust prediction of development to blastocyst stage could significantly simplify and possibly improve the ranking of embryos according to their developmental potential. As shown also by our study, classical morphological criteria applied by an experienced embryologist have a measurable ability to predict development to blastocyst stage. Nevertheless, a higher predictive power is required to make short-term culture competitive with blastocyst culture. An approach similar to that adopted in our study was used by Conaghan et al. (Conaghan et al., 2013). These authors reported that an algorithm applied to TLM-based automatic annotation of embryo morphokinetics improves the ability of experienced embryologists to predict development to the blastocyst stage. The potential of automatic morphokinetic annotation and embryo selection at cleavage stages emerged also in another study based on the same technology (Kieslinger et al., 2016). While these studies had the merit to introduce the concept of combined operator-automated assessment, compared to our data it should be noted that in such cases embryo assessment was extended to day three.

The ability of morphokinetic algorithms generated by TLM technology to predict development to blastocyst and implantation has been extensively investigated as early as 2010 (Wong et al, 2010; Meseguer et al., 2011), although without reaching a consensus (Armstrong et al., 2019). This has prompted the need, recently tackled with the application of AI, to harness the immense potential of TLM to generare morphokinetic data. An initial experience documenting the application of AI to oocyte and embryo scoring in clinical embryology was published as early as 2013 (Manna et al., 2013). Although relevant as a proof-of-principle and based on static images, this study was not followed by more extensive, independent investigations. The first major studies combining TLM technology and AI applied to human IVF were published only last year. Tran et al. (Tran et al., 2019) reported on a modality based on deep learning and raw data derived from TLM sequences of embryos developed to the blastocyst stage. In this research, the authors showed that the probability to achieve a clinical pregnancy can be automatically predicted with an AUC of 0.93. This study however requires independent confirmation, especially because its methodology appears largely unreported. In another study, Khoshravi et al. (Khosravi et al., 2019) demonstrated that an AI platform trained on blastocyst images scored by experienced embryologists can predict blastocyst quality with an AUC >0.98 and the chances of pregnancy in a range between 13.8% and 66.3%, depending on blastocyst and patient characteristics. In addition, Rad and colleagues (Rad et al., 2019) reported that AI can predict embryo implantation based on the analysis of a single blastocyst image. A similar approach was adopted more recently also by VerMilyea and colleagues (VerMilyea et al., 2020), who described an improvement produced by AI of 42% over routine embryologist assessment. These studies, together with others (Curchoe et al, 2019; Dirvanauskas et al., 2019; Kanakasabapathy et al., 2019), have stirred considerable interest in the field and will certainly be followed up by exciting developments.

Combining different technical and analytical tools, we conceived the present study to test the hypothesis that novel information derived from cytoplasmic movements occurring in the early embryo can be harnessed to develop predictive models of development.

Of note, focus on cytoplasmic movements as a source of data to predict human embryo development is a novelty in MAR. In fact, all embryo evaluation models, both static and dynamic, rely on large-scale embryo morphological attributes (e.g. pronuclear position, cell number, degree of fragmentation, rate of blastocele expansion) and times of specific events (cleavages, compaction, start of blastocel formation), leaving cytoplasmic dynamics totally neglected. Therefore, in PIV analysis of cytoplasmic movements interpreted with the power of AI, we glimpsed a novel window of opportunity to improve early prediction of embryo developmental ability. Previous animal studies highlighted the significance of cytoplasmic dynamics for oocyte and embryo function. In a mouse model, Ajduk and colleagues (Ajduk et al., 2011) were the first to demonstrate that TLM associated with PIV can detect specific patterns of cytoplasmic flows occurring during fertilisation, which can be then used to predict embryo viability. Using the same image detection technology, we previously showed that mouse oocyte developmental competence, as assessed by chromatin rearrangement at the germinal vesicle stage, can be correctly predicted with a probability of 90% (Bui et al., 2017). Crucially, however, in this study we made more far-reaching and effective use of the data generated by TLM-PIV, by training an ANN. Therefore, we decided to exploit the excellent predictive power of this approach to assess embryo developmental competence *in vitro* in an MAR scenario. We recognise the limitations of our study, mainly represented by the small data set and the endpoint (blastocyst development), which is not the ultimate goal of infertility treatment. At the same time, however, we are encouraged by the significance and outcome of our study, which however should be intended as a proof-of principle. Firstly, compared with the AI studies of Tran et al. (Tran et al., 2019) and Khoshravi et al. (Khosravi et al., 2019), we used a completely different and new source of data, i.e. analysis of cytoplasmic movement detected by PIV, as discussed above. This demonstrate that non-invasive bright field microscopy can potentially offer more than simply static morphology or morphokinetic information. In particular, our data suggest that sophisticated tracking of subcellular characteristics (discrete cytoplasmic particles) can add a “layer” of morphological information, qualitatively and quantitatively significant, never used or even considered so far for the analysis of human or animal cleavage-stage embryos. Secondly, our data suggest that AI has the potential to improve the predictive ability of an experienced embryologist, achieving 82.6% accuracy, 79.4% sensitivity and 85.7% specificity to predict development to blastocyst. Human intervention may be seen as an obstacle to full automation. On the contrary we believe that valuable human skills should not be lost as an effect of technological development, but rather preserved and used. Thirdly, and particularly significant for the practice of MAR, our study suggests that blastocyst development can be predicted by day two, instead of day three as shown in other studies (Conaghan et al., 2013). Overall, however, two major limitations of our studies should be noted. The small sample size is compatible with the exploratory and preliminary nature of this investigation, but represents a weakness that prevents immediate application in a clinical scenario. Therefore, larger studies should be undertaken to confirm the validity of the present data and assure more stringent control of possible biases derived from clinical parameters. Indeed, we are in the process to set up a larger study focused on clinical endpoints. Not only will this require a larger study population and more participating clinics, but also an evolution of computational and IT tools in order to automate crucial steps of data extraction and elaboration, whose length and manual operativity represent an impediment to routine use of our model.

From a biological perspective, the study confirms the importance of early development for the correct unfolding of later stages. Interestingly, our approach could be further focused to the fertilization window, to assess the specific predictive power of this developmental segment. Consistent with this proposition, it is interesting to note that PIV patterns of competent and non-competent embryos diverge significantly during the interval of fertilisation comprised between pronuclear positioning and breakdown. Once again, this highlights the importance and predictive power of fertilisation events, as reported in recently published studies (Coticchio et al., 2018). Alternatively, the present approach could be extended to the cytoplasmic activity of human unfertilized oocytes, mirroring our previous mouse study (Bui et al., 2017).

From a more practical standpoint, should these preliminary data be confirmed, the possibility to limit embryo culture to two days would entail a reduction in costs and complexity of the IVF process compared with day five culture, while preserving a clinical efficiency comparable with blastocyst transfer.

## Conclusions

Collectively, although not immediately translatable into clinical embryology practice, the present study suggests that movement of cytoplasmic particles of the human early embryo is a novel and valuable source of data to predict further development, after elaboration by advanced imaged analysis and AI. This opens new perspectives for non-invasive embryo assessment. Indeed, we are in the process to gather forces for a much larger study having live birth rate, as primary endpoint, and other parameters of clinical outcome as secondary endpoints.

## Acknowledgments

This work was made possible thanks to support of 9.baby Family and Fertility Center, the Italian Ministry of Education, University and Research (MIUR) Dipartimenti di Eccellenza Program (2018–2022) to the Department of Biology and Biotechnology ‘L. Spallanzani’, University of Pavia, and a grant from the University of Pavia (FRG 2018). The authors also thank Merck-MilliQ Laboratory Water Solutions for support.

